# Applications of a Novel Reciprocating Positive Displacement Pump in the Simulation of Pulsatile Arterial Blood Flow

**DOI:** 10.1101/2022.06.20.496802

**Authors:** Adam Menkara, Ahmad Faryami, Daniel Viar, Carolyn A Harris

## Abstract

Pulsatile arterial blood flow plays an important role in vascular system mechanobiology, especially in the study of mechanisms of pathology. Limitations in cost, setup time, sample size, and control across current in-vitro and in-vivo methods prevent future exploration of novel treatments. Presented is the verification of a novel reciprocating positive displacement pump aimed at resolving these issues through the simulation of human ocular, human fingertip and skin surface, human cerebral, and rodent spleen organ systems. A range of pulsatile amplitudes, frequencies, and flow rates were simulated using pumps made of 3D printed parts incorporating a tubing system, check valve and proprietary software. Volumetric analysis of 430 total readings across a flow range of 0.025ml/min to 16ml/min determined that the pump had a mean absolute error and mean relative error of 0.041 ml/min and 1.385%, respectively. Linear regression of flow rate ranges yielded R^2^ between 0.9987 and 0.9998. Waveform analysis indicated that the pump could recreate accurate beat frequency for flow ranges above 0.06ml/min at 70BPM. The verification of accurate pump output opens avenues for the development of novel long-term in-vitro benchtop models capable of looking at fluid flow scenarios previously unfeasible, including low volume-high shear rate pulsatile flow.

## Background

Pulsatile arterial blood flow has been shown to have inherent properties that are integral to the normal function of specific organ and tissue systems[1,2]. The ability to generate pulsatile flow is important in the development of in-vitro models to simulate real-life conditions. However, many modern fluidic devices resort to utilizing constant or harmonic-based fluid flow. This is especially true in low volume conditions due to limitations in producing physiologically accurate low volume flow[3].

While extremely useful when studying systemic impact, animal studies are limited in their ability to directly and simultaneously mimic human physiologic arterial blood flow. These studies are also usually very time-intensive to setup and are limited in the range of physiologic and pathophysiologic scenarios that can be measured and tested dynamically. In-vitro models maintain control and dynamic recording potential. However, significant cost and time investment required to build, verify, and validate a test setup prior to conducting flow based biomechanistic in-vitro studies limits their usage.

This study presents the testing and verification of a novel positive displacement pump and operating program through the simulation of literature-based blood flow data of organ systems spanning human ocular, human fingertip and skin surface, human cerebral, and rodent spleen. These blood flow data span arbitrarily defined low (0ml/min – 0.4ml/min), mid (0.4ml/min – 1.3ml/min) and high (3ml/min – 16ml/min) flow rate ranges aimed to provide flow rate performance for different flow applications. The use of a syringe-based positive displacement method of driving fluid flow in conjunction with the precise control of programmable input variables through a user interface allow for the generation of a wide-range of pulsatile low volume fluid flow rates and waveforms. Furthermore, the incorporation of a robust check valve system enables the ability for the pump to automatically reset back to its original position allowing the pump to function in long term fluid flow experiments.

## Materials and Methods

### Reciprocating Positive Displacement Pump

The body of each reciprocating positive displacement pump is comprised of 3D printed parts made of polylactic acid plastic (PLA) manufactured using an Anycubic I3 Mega printer (Anycubic Technology CO., Limited, HongKong)[4]. The design consists of five 3ml syringes per pump, allowing flow output from five total channels per pump (Fig 1.). A stepper motor, linear bearings, motor coupler, and lead screw assembled in line with each channel allow for the conversion of rotational motion into linear movement.

**Fig 1.**
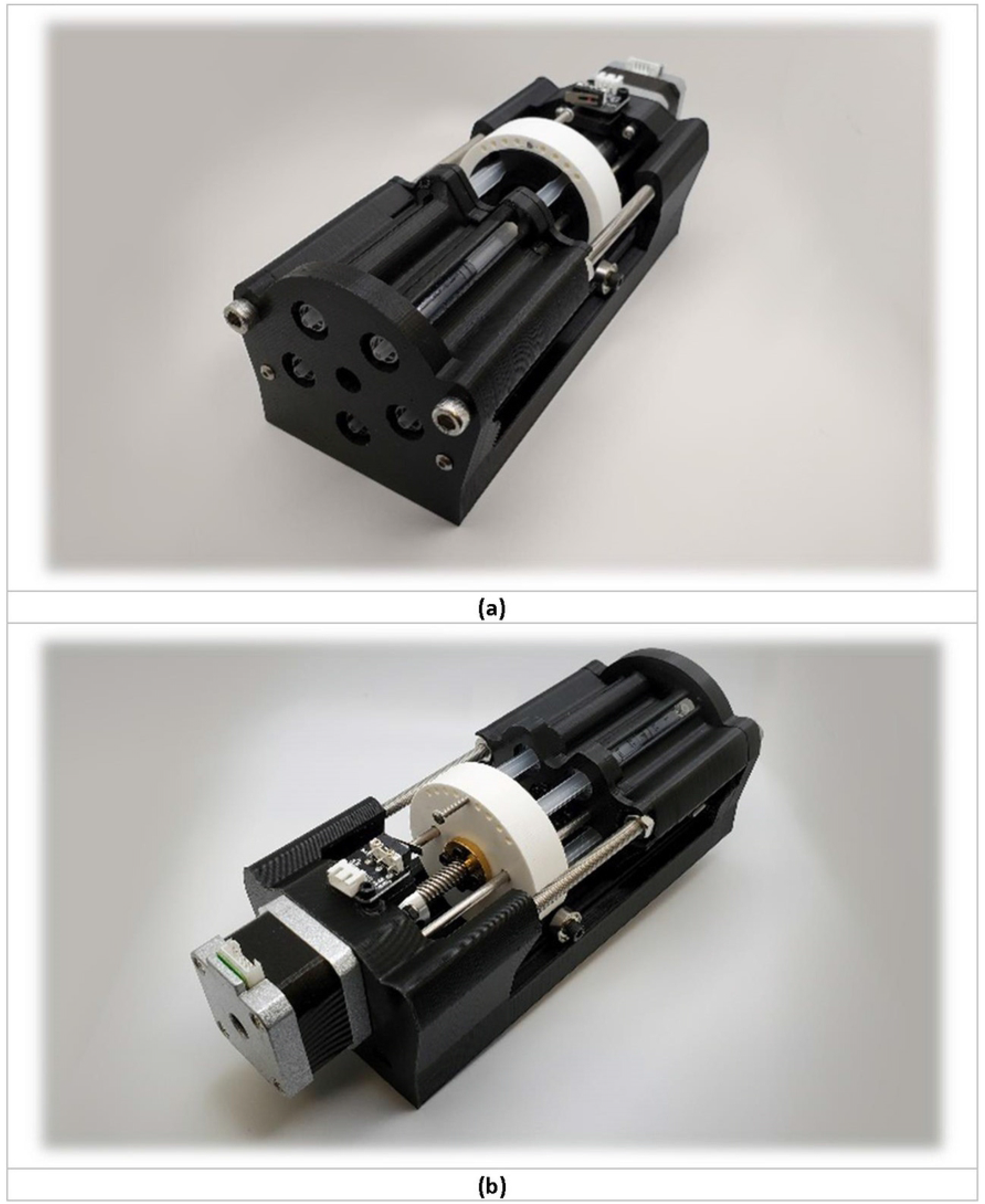
Reciprocating Positive Displacement Pump. Front view (a) and rear view (b) of fully assembled reciprocating positive displacement pump

A proprietary software allowing for the direct input through a simple user interface of user profile variables consisting of output bulk volume rate, pulsation frequency, and pulsation amplitude was developed using Python. Based on user input per profile, rotational speed and duration were calculated by the software. Mechanical inefficiencies of the pump and other components were account for through the program. This information is then sent to an Arduino-based board which is then interpreted and used to drive stepper motor movement. Each Arduino-based board is capable of simultaneously running three pumps, a total of 15 channels per board. An automatic sterilization function was also built within the program consisting of a constant reciprocating action of the pump for a total of 20 minutes, cycling 99% isopropyl alcohol throughout the duration of the cleaning function. This is followed by three cycles of deionized water to expel any remaining isopropyl alcohol from the tubing system.

Once the syringes have reached their fully compressed state, the program then automatically resets the pump back to its original position, extending the syringes fully. The use of individual check valves immediately after the output from the syringes enables this retracting motion of the pump to automatically refill the syringes. This retraction time takes a total of approximately eight seconds. Each check valve limits the chances of cross-contamination between sample outputs and between input and output within each channel. Luer locking fittings used by the check valves and syringes allowed for simple and robust fluidic connections between all components. Input and output for each individual channel consisted of 50cm 3mm inner diameter silicone tubing into their respective check valves.

### Systolic Time Interval

An important variable in the accuracy of individual pulsations is systolic time interval. An analogous representation of ventricular systolic time interval is created through the implementation of the systole time user input variable. This allows for direct control over beat cycle length independent of volume rate or frequency. Ideally, beat cycle duration can be approximately equal to two times the input systole time, although external variables such as system compliance may affect this.

### Amplitude

Amplitude of individual pulsations is based on the multivariable user input per profile. Amplitude is directly correlated to volume rate and compliance of the tubing and flow system, and inversely correlated to both systole time and beat frequency. The dampening factor is a representation of compliance of the entire tubing system, and refers to the tubing system itself, attached chambers to the tubing and fluid viscosity. All these variables such as tubing diameter can be manipulated to better fit the intended modelling parameters. The full relationship between these variables and their effects on amplitude can be seen in Equation 1:

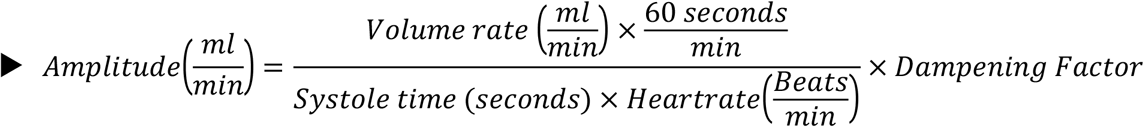

The incorporation of a method of simply and directly manipulating pulsatile amplitude allows for the freedom to explore domains of clinical applications previously unexplored.

### Testing Parameters

In our previous study, the accuracy of the reciprocating positive displacement pump was demonstrated across a range of 0.01ml/min to 0.7ml/min with an R^2^ value of 0.9998[5]. To expand the verified flow rate range and investigate the capabilities of the pump in novel applications of arterial blood flow in organ systems, a literature search was conducted collecting flow rate data spanning three ranges; low (0ml/min – 0.4ml/min), mid (0.4ml/min – 1.5ml/min) and high (3ml/min – 16ml/min) bulk flow rates. The organ systems used to populate these arterial blood flow ranges are human ocular, human fingertip, human cerebral, and rodent spleen (Table 1). Unless otherwise noted, an input of 70BPM and 0.1 second systolic time were used for all human simulations and 380BPM and 0.054 second systolic time for all rat-based simulations[6–8]. While any heartrate within physiologic domain could have been selected, an average resting 70BPM for humans and 380BPM for rats was chosen unless otherwise specified. Systolic time was not provided in any study simulated throughout the flow ranges, therefore requiring the use of arbitrary systolic times of 0.1 seconds for human and 0.054 seconds for rat-based simulations.

**Table 1.** Organ systems scenarios and corresponding flow rate, beat rate, and systolic time input profiles into reciprocating positive displacement pump.

| Input Flow Profile for each organ system, measuring device and condition | Flow Rate (ml/min) | Beat Rate (BPM) | Systolic Time (seconds) | Reference |
| --- | --- | --- | --- | --- |
| **Low Flow Range (0ml/min – 0.4ml/min)** |  |  |  |  |
| Fingertip blood flow autoregulation (post-caffeine) | 0.029±0.004 | 71 | 0.1 | [9] |
| Retinal Blood Flow by Laser Doppler Velocimetry | 0.033 | 70 | 0.1 | [10,11] |
| Fingertip Blood Flow by Venous-occlusion volume plethysmography (lower) | 0.056 | 70 | 0.1 | [12] |
| **Fingertip blood flow autoregulation (pre-caffeine)** | 0.067±0.009 | 74 | 0.1 | [9] |
| **Retinal Blood Flow by Laser Doppler Velocimetry** | 0.08 ±0.012 | 70 | 0.1 | [13] |
| **Fingertip blood flow autoregulation (baseline)** | 0.088±0.015 | 76 | 0.1 | [9] |
| **Fingertip Blood Flow by Venous-occlusion volume plethysmography (Critical vasoconstriction Temperature)** | 0.2 | 70 | 0.1 | [12] |
| **Retinal Blood Flow by Phase Contrast MRI** | 0.261±0.087 | 70 | 0.1 | [14] |
| **Mid Flow Range (0.4ml/min – 1.4ml/min)** |  |  |  |  |
| **Fingertip Blood Flow by Venous-occlusion volume plethysmography** | 0.42 | 70 | 0.1 | [12] |
| **Total Pulsatile Ocular Blood Flow by MRI (lower)** | 0.444 | 70 | 0.1 | [14] |
| **Rat Splenic Arteriovenous Flow Differential in Rats (*pre-caudal ligation*)** | 0.6±0.1 | 380 | 0.054 | [15] |
| **Splenic Arteriovenous Flow Differential in Rats (*pre-caudal ligation*)** | 0.8±0.3 | 380 | 0.054 | [15] |
| **Total Pulsatile Ocular Blood Flow by MRI (upper)** | 0.803 | 70 | 0.1 | [14] |
| **Ocular Choroidal Blood Flow by MRI** | 0.917±0.281 | 70 | 0.1 | [14] |
| **Total Pulsatile Ocular Blood Flow by Langham Pneumotonometer** | 1 | 70 | 0.1 | [16,17] |
| **Splenic Arteriovenous Flow Differential in Rats (*post-caudal ligation*)** | 1.0±0.2 | 380 | 0.054 | [15] |
| **Splenic Arteriovenous Flow Differential in Rats (*post-rostral ligation*)** | 1.2±0.1 | 380 | 0.054 | [15] |
| **High Flow Range (3ml/min – 16ml/min)** |  |  |  |  |
| **Middle Meningeal Artery** | 6±3 | 70 | 0.1 | [18] |
| **Ophthalmic Artery** | 11±5 | 70 | 0.1 | [18] |

These systolic times are used as a representation of true systolic time, however changes in systolic times can be made for direct manipulation of pulsatile amplitude independent of flow volume rate and beat rate. Degassed, deionized water, measured and verified at 1 gram/milliliter of water at room temperature, was used to measure the accuracy and precision of pump output. A total of 10, ten-minute weight measurements were performed across 10 pump channels using the pump and tubing setup schematic seen in Fig 2. per input user profile. The sterilization function described earlier was used to properly clean and prime pumps before each testing session. Individual beakers were used per channel, weighed after the time period of each test was completed. Weight measurements were conducted using a Mettler Toledo AT261 DeltaRange Analytical Balance (Mettler-Toledo, LLC, USA).

**Fig 2.**
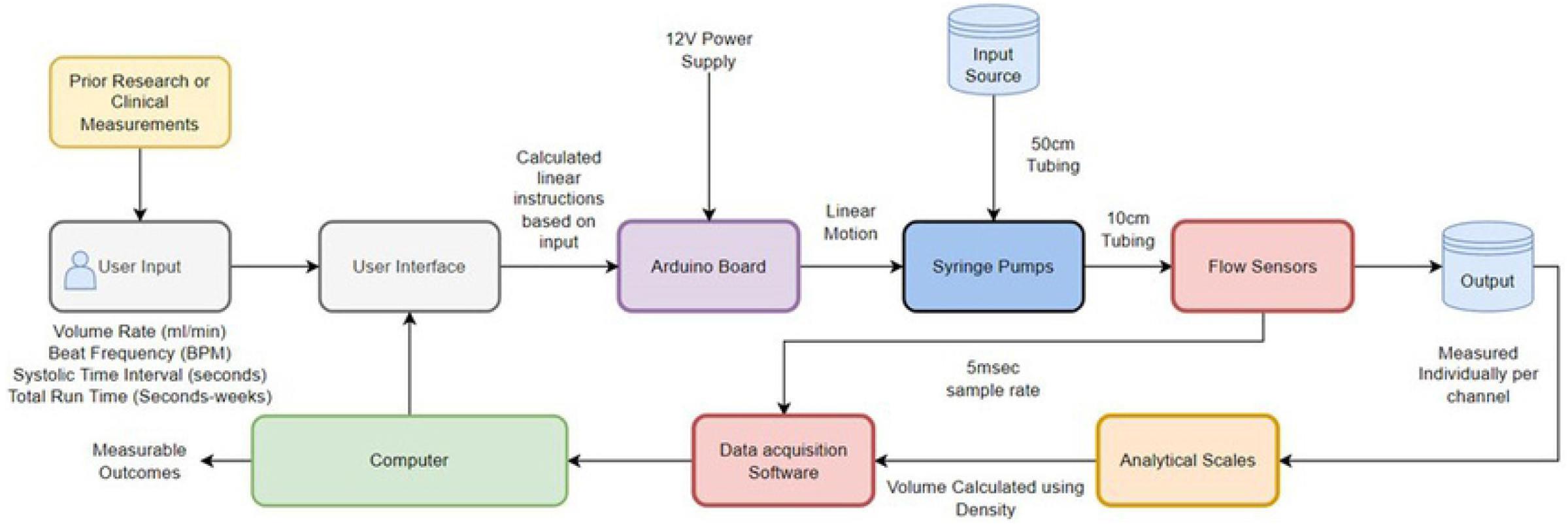
Schematic diagram of testing setup. Illustrates flow of fluids and data collection locations of user input verification

Each individual profile tested also had flow data obtained from a flow sensor placed at the immediate output of a channel check valve for each individual profile tested. The flow sensor used to collect flow data was Sensirion SLF06 series flow sensor (Sensirion AG, Switzerland). Flow data was used to determine consistency of amplitude and volume rate within the same test, accurate beat rate throughout the duration of each test, and differences of volume rate and amplitude across different profiles.

## Results

### Volumetric Analysis

The pump was able to simulate the volumetric flow rates for the low, mid, and high bulk flow rate ranges spanning a total range of 0.025ml/min to 16ml/min for the organ systems of the human eye, human fingertip and skin surface, rat spleen and human cerebral blood flow rates (Fig 3, Table 2).

**Fig 3.**
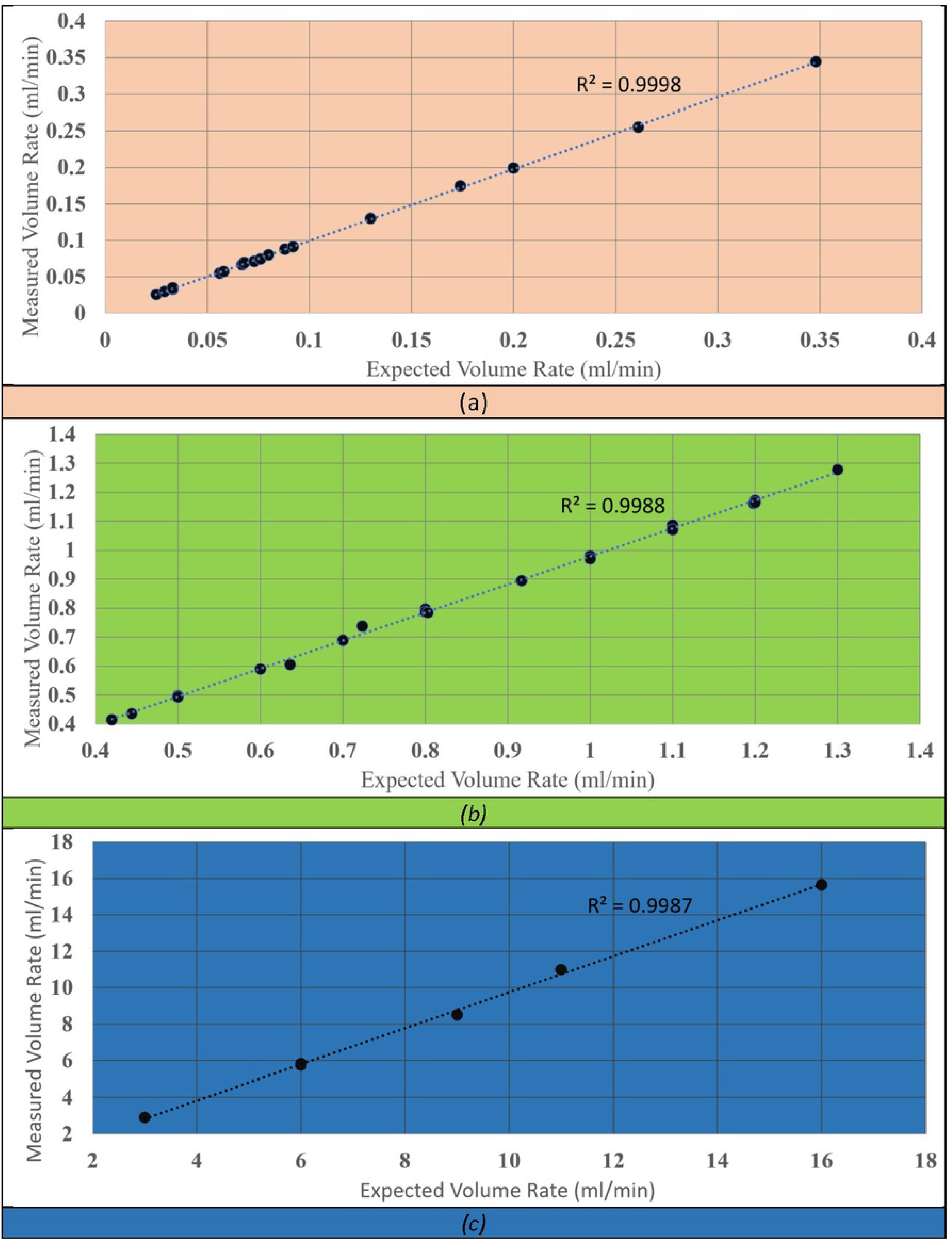
Expected vs. Measured low, mid, and high bulk flow rate ranges. Illustrates the expected versus mean of measured volume rate across 10 channels for low flow ranges (0ml/min – 0.4ml/min) (a), mid flow ranges (0.4ml/min – 1.4ml/min) (b), and high flow ranges (3ml/min – 16ml/min) (c) with a linear function showing a regression and R^2^ value for each plot

**Table 2.** Verified organ systems explored in this paper within achievable range of reciprocating positive displacement pump.

| Organ System | Minimum Input Flow Rate ($\frac{ml}{min}$) | Maximum Input Flow Rate ($\frac{ml}{min}$) | Pump Achieves Physiologic Flow |
| --- | --- | --- | --- |
| Retinal Blood Flow | 0.033 | 0.348 | Yes |
| Ocular Choroidal Blood Flow | 0.636 | 1.198 | Yes |
| Total Pulsatile Ocular Blood Flow | 0.444 | 1 | Yes |
| Fingertip Blood Flow | 0.025 | 0.42 | Yes |
| Splenic Arteriovenous Flow Differential in Rats | 0.5 | 1.3 | Yes |
| Ophthalmic Artery | 6 | 16 | Yes |
| Middle Meningeal Artery | 3 | 9 | Yes |

Across the entire range of flow rates, a total of 430 ten-minute weight measurements taken from the pumps had a mean absolute error and mean relative error of 0.040854 ml/min and 1.385164% respectively. The linear regression of the low, mid, and high flow rate ranges (Fig 3, a-c, respectively) tested yielded R^2^ values 0.9998, 0.9988 and 0.9987, respectively. The standard deviation across the total range of 0.025ml/min to 16ml/min was 0.00151ml/min to 0.14196ml/min presented in Fig 4(a). Relative standard deviation indicates the pumps’ precision across the range of tested flow rates (Fig 4(b)). A novel verified range for pump input parameters can be seen in Table 3.

**Fig 4.**
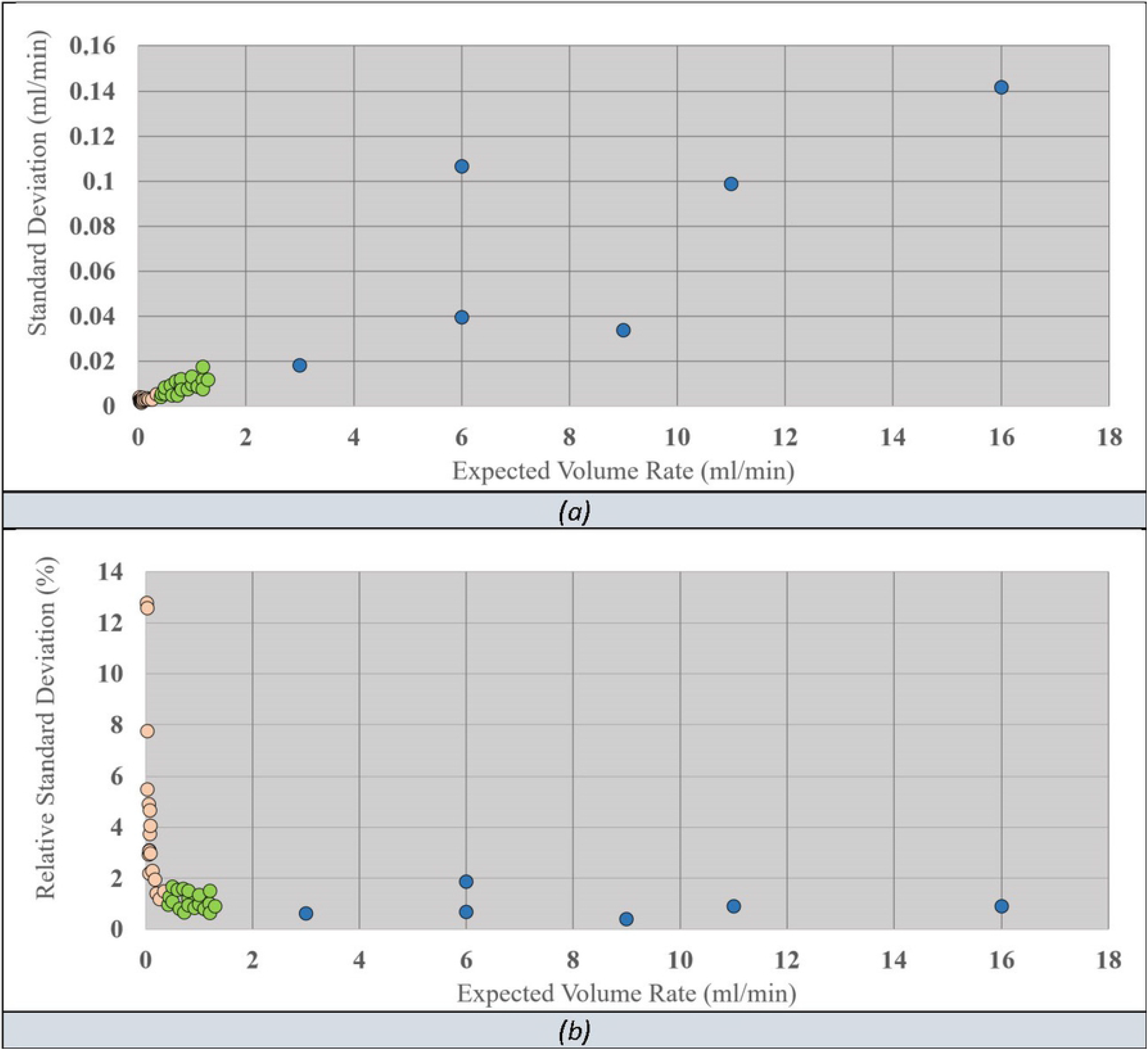
Standard Deviation across full bulk flow rate range. Illustrates the standard deviation(a) and relative standard deviation(b) across full range of tested flow ranges

**Table 3.** Verified pulsation rate, volume rate and amplitude range achievable by the reciprocating positive displacement pump.

| Category | Minimum | Maximum | Capable of Achieving? |
| --- | --- | --- | --- |
| Pulsation Rate ($\frac{Beats}{min}$) | 1 | 400 | Yes |
| Output Volume Rate ($\frac{ml}{min}$) | 0.01 | 16 | Yes |
| Amplitude ($\frac{ml}{min}$) | 0.02 | 65 | Yes |

### Pulsatile Flow

Flow measurements from the pump illustrated accurate pulsatile flow according to beat rate and volume rate in humans and rats (Fig 5-7). Flow rates under 0.06ml/min with an input of 70BPM were too small and were unable to simulate accurate 70BPM pulsatile waveform flow.

**Fig 5.**
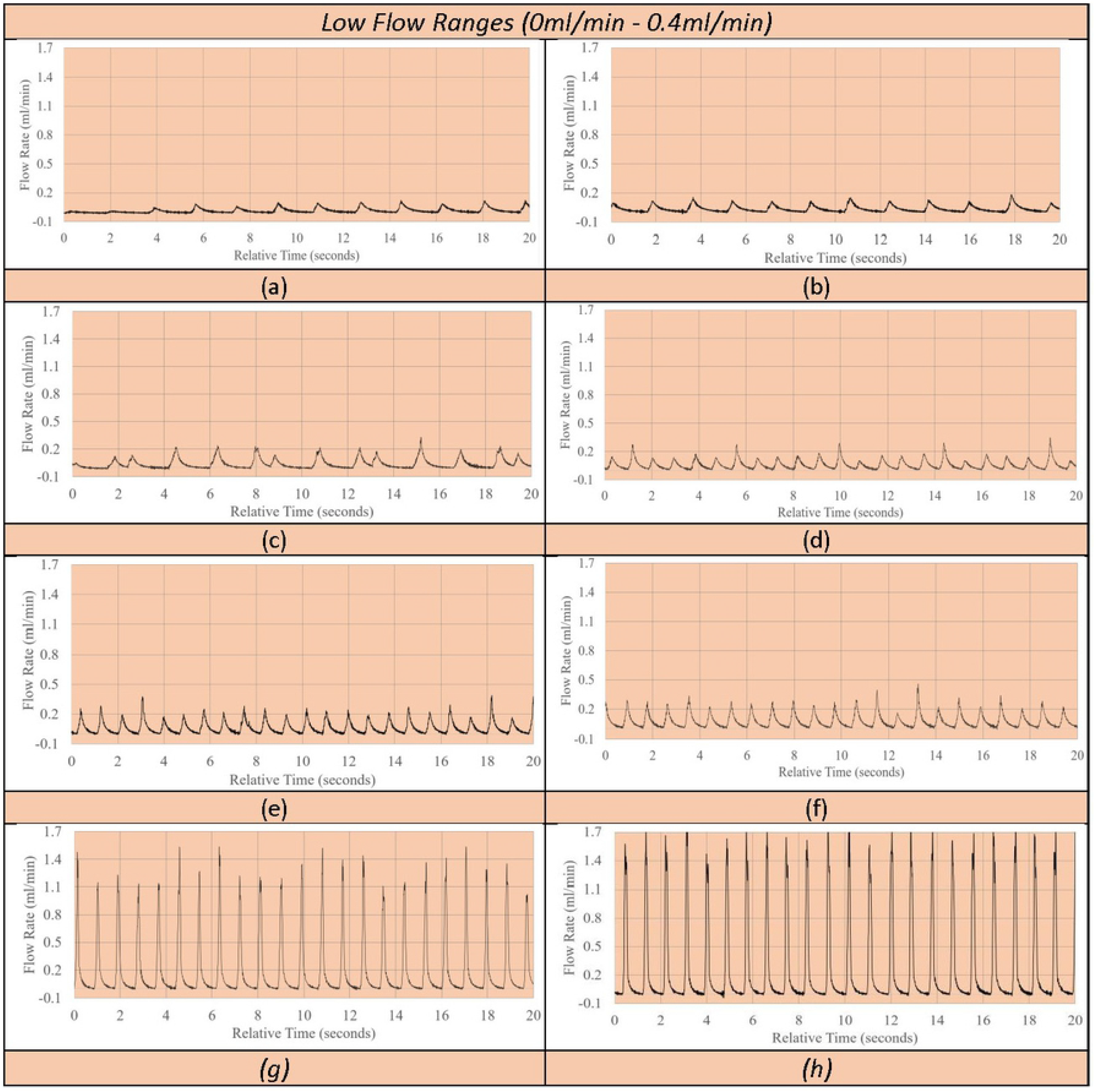
Low flow range pulsatile waveform patterns. Visual representation of flow waveform pattern of mean flow in order from lowest to highest bulk volume rate simulated in low flow rate range; effects of caffeine on fingertip blood flow autoregulation (post-caffeine) (a), retinal blood flow by laser doppler velocimetry (b), fingertip blood flow by venous occlusion plethysmography (c), effects of caffeine on fingertip blood flow autoregulation (pre-caffeine) (d), retinal blood flow by laser doppler velocimetry (e), effects of caffeine on fingertip blood flow autoregulation (baseline) (f), critical vasoconstriction temperature for fingertip blood flow (g), retinal blood flow by phase contrast MRI (h)

**Fig 6.**
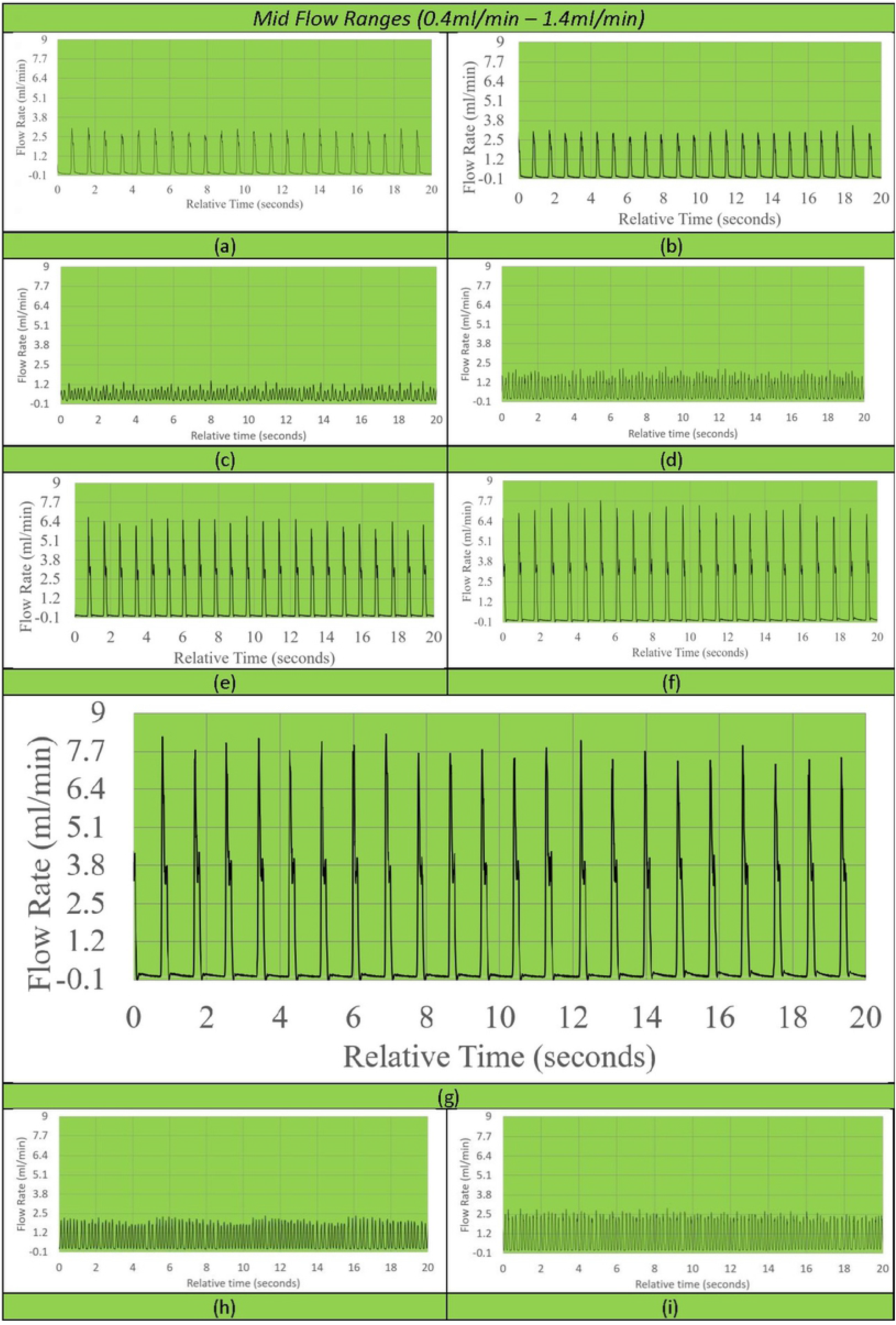
Mid flow range pulsatile waveform patterns. Visual representation of flow waveform pattern of mean flow in order from lowest to highest bulk volume rate simulated in mid flow rate range; fingertip blood flow by venous occlusion plethysmography (a), total pulsatile ocular blood flow by phase contrast MRI (b), splenic arteriovenous flow differential in rats (pre-caudal ligation) (c), splenic arteriovenous flow differential in rats (pre-rostral ligation) (d), total pulsatile ocular blood flow by phase contrast MRI (e), ocular choroidal blood flow by phase contrast MRI (f), total pulsatile ocular blood flow by Langham pneumotonometer (g), splenic arteriovenous flow differential in rats (post-caudal ligation) (h), splenic arteriovenous flow differential in rats (post-rostral ligation) (i)

**Fig 7.**
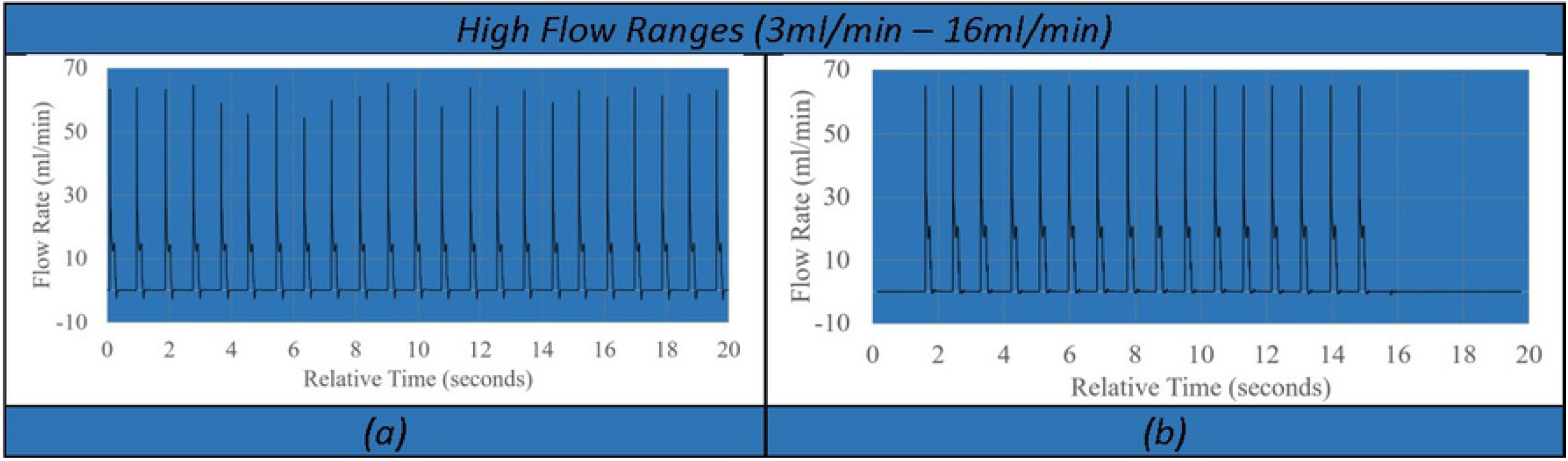
High flow range pulsatile waveform patterns. Visual representation of flow waveform pattern of mean flow in order from lowest to highest bulk volume rate simulated in high flow rate range; Middle meningeal Artery (a), Ophthalmic Artery (b)

Therefore, waveforms presented in Fig 5 (a-c) do not have the correct beat counts in the relative time frames, although the pump remains volumetrically correct due to inbuilt catchup functions within the proprietary program. However, A total of ~23 total beats were counted across the 20 seconds displayed for all other displayed flow patterns, equating to the input value of ~70BPM.

To better illustrate the effects of input volume changes on individual pulsation amplitude and pattern, Fig 8 (a-d) illustrates a 1 second, single beat comparison of mean, upper and lower flow rate limits of standard deviation of measurement in ocular choroidal, total pulsatile ocular, and retinal blood flow rates. One second, single beat comparisons were not shown for organ system simulations where flow rates dropped below 0.06ml/min, waveform amplitudes exceeded the flow rate limit of the flow sensor used (65ml/min), or similarity in single beat waveforms made the subsequent graph unclear. Fig 9 illustrates the direct simulation done using the reciprocating pump of retinal blood flow rates patterns from previous literature, matching fluid rise times, peak amplitude, and total beat time similar to the Figure presented by Rebhan et. al[19].

**Fig 8.**
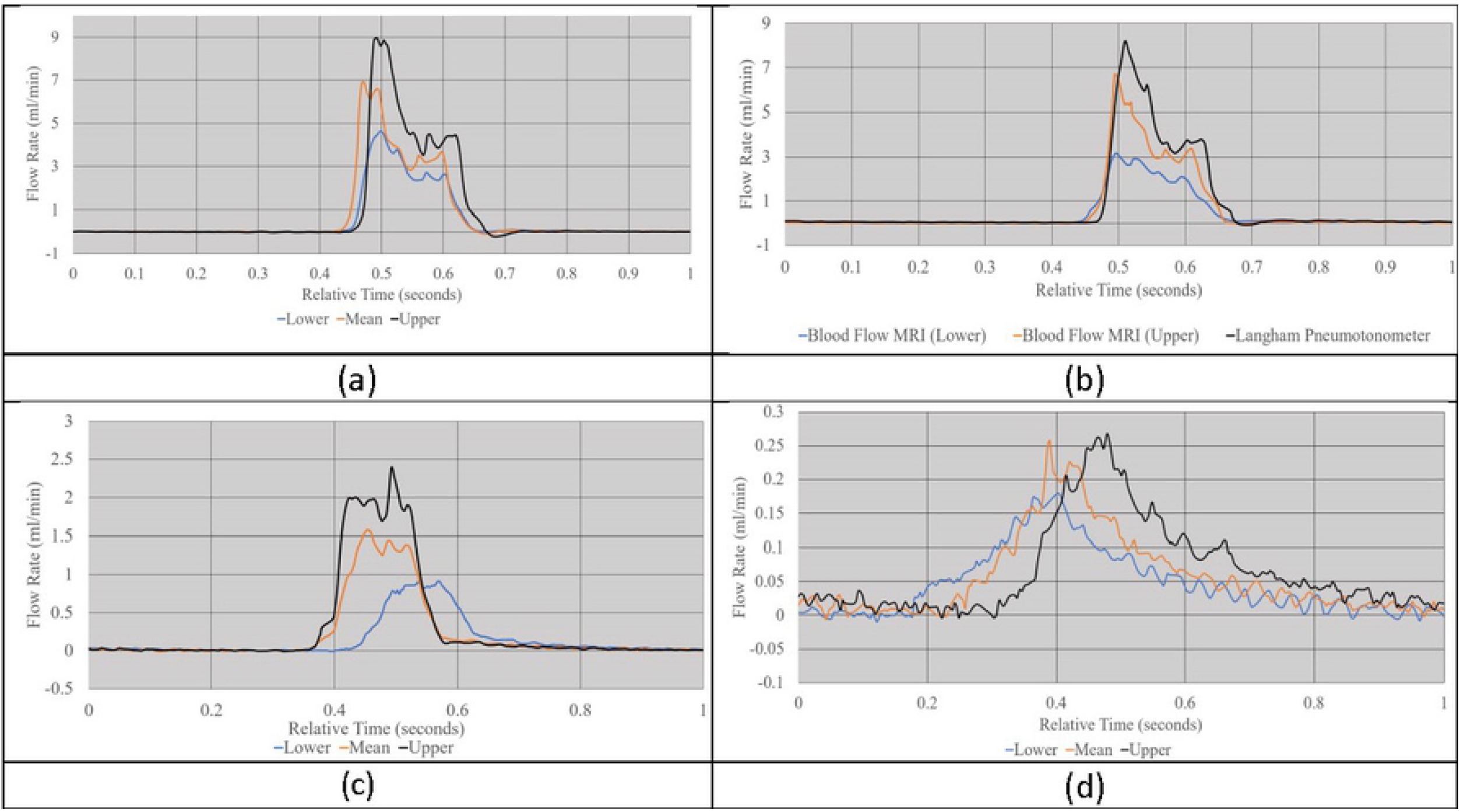
Visualization of single second pulsatile waveform patterns. Visual Representation of comparison of lower, mean, and upper limits of flow waveform patterns across 1 second for choroidal blood flow rates (a), total pulsatile ocular blood flow measurements (b), retinal blood flow gathered through MRI (c), retinal blood flow gathered using Laser Doppler velocimetry (d)

**Fig 9.**
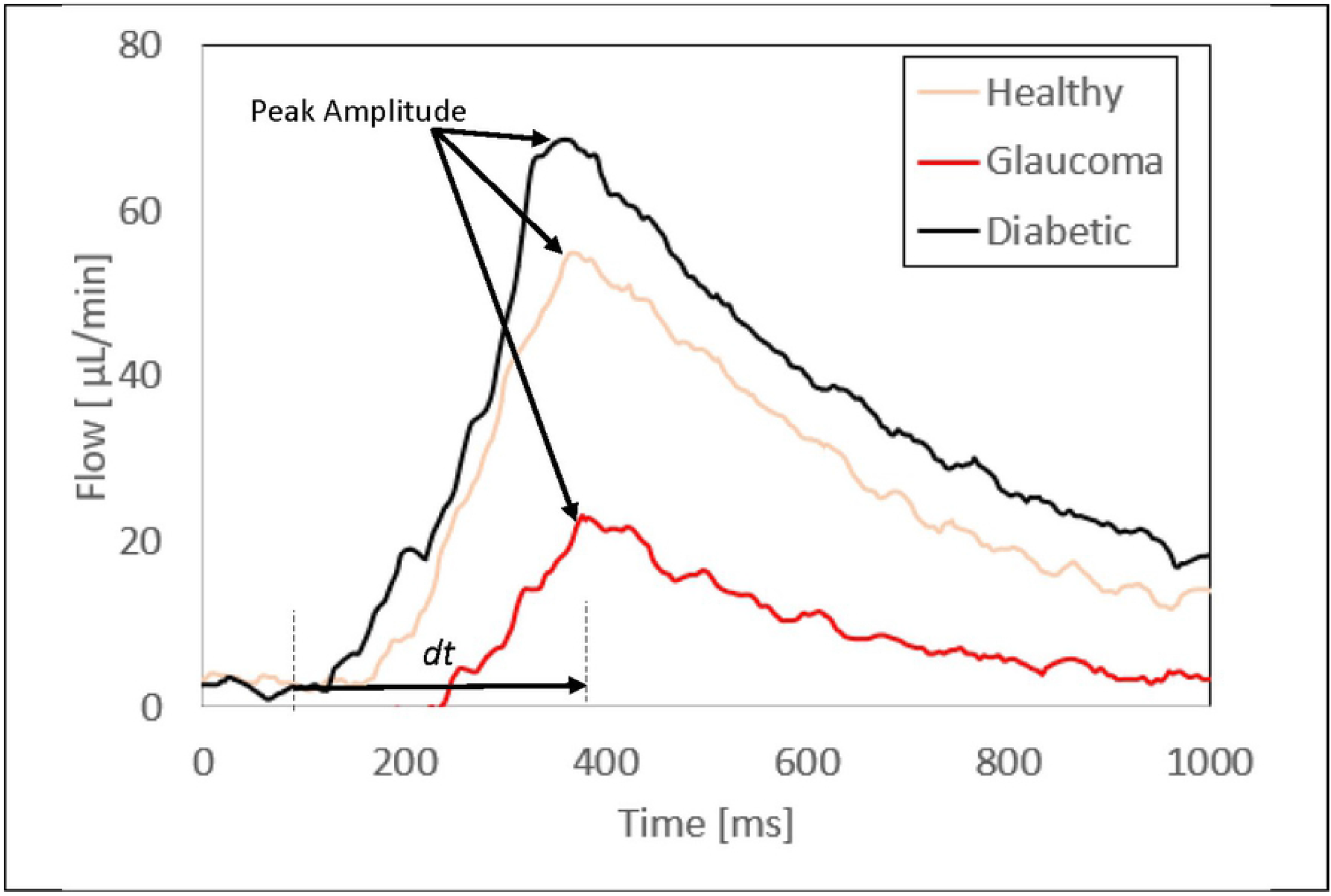
Waveform pattern simulation. Visual representation of simulation using reciprocating pump of one-second flow waveform patterns of retinal blood flow in a healthy, diabetic, and glaucoma patient scenarios generated from a computational framework

In the simulation seen in Fig 7 (b), very high flow rates (above 6ml/min) were high enough to require the use of the reciprocating action of the pump before the full 20 seconds has elapsed, requiring the pump to return to its original, fully retracted position state and explains the lack of pulsatile flow across the full 20 seconds.

## Discussion

Using the testing setup, volumetric analysis of pump output was accurate within a standard deviation range of 0.00151ml/min to 0.14196ml/min through the low (0ml/min – 0.4ml/min), mid (0.4ml/min – 1.3ml/min) and high (3ml/min – 16ml/min) bulk flow rate ranges within the organ systems tested (Fig 3). Through this, the pump has shown to provide accurate flow in ranges in which the development of mechanistic in-vitro models and products requires. Although specific organ systems were used to define the flow ranges tested, these simulations for organ systems are used as example vascular bodies and the versatility of the pump expands beyond these examples. This versatility is also created through the implementation of the reciprocating action of the pump once syringe limits have been reached. The reciprocating action of the pump takes approximately eight seconds and stops flow for that duration of time. However, the total bulk flow output, beat frequency and time input of flow are not affected.

This is one way in which the reciprocating positive displacement pump system distinguishes itself, making it ideal for use in long term fluid modeling scenarios where output variables from the model are dependent on input pulsatile flow. The use of the proprietary program along with its ability to take in user input for heart rate, volume rate and amplitude, also employs a user-input time variable where the program can calculate the exact number of pulsations within the input time frame and executing them, excluding time of retraction. This in turn, allows for precision in long term applications of the pump and makes the versality of pump applications span much wider than the organ systems described throughout this paper, although the applications within the organ systems are vast in themselves.

### Human Ocular and Retinal Blood Flow

Unlike previous pump flow systems, non-harmonic pulsatile flow can be achieved, through the manipulation of volume rate, beat rate and amplitude. This means that low volume pulsatile flow simulations are no longer limited by the method of simulating them, but rather the accuracy of the measurement method itself and its application, assuming it is within the achievable range of the pump. This can be seen through the successful simulation of multiple methods of measuring retinal and total pulsatile ocular blood flow seen in Table 1 and Fig 3. Individual pulsations utilizing our reciprocating pump are also capable of being simulated at low flow rates, through the manipulation of BPM and amplitude input. This allowed the pump to reproduce the individual pulsations of blood flow through the human retina seen in Fig 9. Previous literature has shown that flow shear stress is an important indicator of vascular disease[20,21]. Our pump, with its ability to directly manipulate flow amplitude, a key component to shear stress in arteries, provides an avenue in which its incorporation into a flow based in-vitro model can provide comprehensive insight into the role shear stress plays in retinal disorders. The use of this system in conjunction with an integrated in-vitro model of the human eye does not need to be confined within the domain of the retina. The successful simulation of choroidal blood flow rates and waveforms makes the pump ideal to be integrated into an in-vitro model, an example an in-vitro investigation of retinal detachment. Retinal detachment has been shown to be correlated to central ocular choroidal blood flow with atrophy of the retinal pigment epithelium[22]. Because of current limitations in the simulation of low volume fluid flow models, static in-vitro models with short perfusion times are often resorted to when looking at this phenomenon[23]. The pumps implementation in in-vitro models can provide a more comprehensive understanding of the effects of blood flow on retinal detachment and other diseases. Innovations in vascular development and perfusion methods for future in-vitro models can further increase model accuracy and applicability of the pump. One early possibility can be seen through the development of optical vascular structures developed using optical coherence tomography[24]. The pulsatile nature of the fluid flow has yet to be accounted for in an in-vitro setup involving the human eye, although waveform dynamics have been shown to affect patient outcome in specific scenarios[25].

### Human Fingertip and Skin Surface Blood Flow

The reciprocating positive displacement pump is also capable of simulating fingertip and skin surface blood flow under varying physiologic and pathophysiologic scenarios. An example application within this specific vascular domain is in the bioprinting of skin grafts for burn victims, where functional vascular endothelial cells require perfusion with a peristaltic pump[26]. The use of peristaltic pumps has been compared to harmonic wave flow because of its low peak-to-peak amplitude per pulse and lack of a significant flow drop between pulsations[27]. Pulsatile shear rate has been shown to increase the proliferation of vascular endothelial cells, meaning the implementation of the reciprocating positive displacement pump in a cell perfusion setup used for 3D printed cells can perhaps improve the rate and complexity at which vasculature develops, and in turn accelerate the progression towards a viable bio-printed skin-graft[23,28]. Not only can the increased shear rates of output fluid flow affect the rate at which vascular tissue develops, but also the contents of the fluids used to perfuse the cellular structure. In our setup, the simple silicone tubing system used alongside the reciprocating positive-displacement pump allows for a vast array of fluids with physiologic or pathophysiologic makeup. An example fluid explored in the previous literature is cerebrospinal fluid (CSF), where the reciprocating positive displacement pump can be used to model pulsatile flow, shear rate, shear stress, and amplitude simultaneous to variations in CSF composition as this is directly relevant to pathologic states like those in hydrocephalus[5,29].

### Splenic Blood Flow in Rats

Non-human simulations using our pump are presented because of possible limitations in the collection of flow data in humans in-vivo and in turn, limitations of data for a range of organ systems. Animal models are the main alternative to in-vitro modeling; although useful in their ability to model systemic effects and device compatibility, are time-intensive to setup, limited in the range of conditions testable, and whose flow conditions may not map to humans linearly. Modeling animal flow rates gives us flexibility to eliminate unknown variables, and perhaps may reflect on human data if there are anatomical similarities between species. An example of this within this organ system domain are splenic baroreceptors and their control over splenic afferent nerve activity[30]. Novel developments in carbon-based organic semiconductors can be implemented as an afferent nerve substitute where their efficacy relative to the original nerve in a rat-based in-vitro model with pulsatile fluid flow can be modeled by the reciprocating positive displacement pump[31].

### Human Cerebral Arterial Blood Flow

The ability to isolate flow from different sources provides further versatility of the reciprocating positive displacement pump, as seen in the ophthalmic artery simulation as well as the total pulsatile ocular and retinal blood flow simulations presented in Table 1 and Fig 3. The use of anticancer chemotherapy drugs has been shown to have complications with many organs in the human body, one being the induction of ocular toxicity and other complications with the eye and retina[32,33]. Past in vitro models of the human eye used primary human retinal endothelial cells, employing relatively short perfusion times of chemotherapeutic agents[34]. The pump can act as a method to perfuse the cells over a much longer period and can be a more comprehensive in-vitro test with more accurate flow shear rates throughout treatment. The isolated effects of a chemotherapeutic drug can also be measured, through the pumps ability to isolate specific flow rates of different vascular systems and arteries and get a much more tailored flow modelling setup. The simulation of the middle meningeal artery has fewer applications for the simulation in and of itself, although this additional application confirms that the pump is capable of simulating flow rates necessary for the study of disease not well implemented to date.

### Limitations of the Reciprocating Positive Displacement Pump

The pump is required to retract back to its initial state and reset after a set time simulating pulsatile fluid flow depending on the volume flow rate input profile used. This retraction time, and in turn pause time on the induction of fluid flow, lasts around eight seconds in the current setup. Although this is a minimal pause in fluid flow production, future iterations of the pump and proprietary program design can significantly reduce this time to near zero ensuring a constant pulsatile fluid flow is delivered to a fluid setup or model being investigated. The current syringe volume of 3ml in the pump limits the range of higher flow rate models that can be simulated, something a peristaltic pump can achieve through the sequential addition of peristaltic pump outputs in series. This can be addressed in future iterations of the pump by increasing syringe volume or using the same approach as a sequential peristaltic pump system where combining channels of the same pump and channels from multiple pumps would deliver a significantly larger volume of pulsatile fluid flow.

### Conclusion

In conclusion, the verification of the reciprocating positive displacement pump determined it was accurate in the simulation of arterial blood flow in human ocular, human fingertip and skin surface, human cerebral, and rodent spleen organ systems. This provides new freedom in the development of novel in-vitro benchtop models involving pulsatile fluid flow and can accelerate the development of translatable treatments to improve patient outcome.

## Declarations

The authors have no conflicts to disclose.

## References

1. Rocha FG. Liver blood flow. Blumgart’s Surgery of the Liver, Pancreas and Biliary Tract. Elsevier; 2012. pp. 74–86.e5. doi:10.1016/B978-1-4377-1454-8.00004-7

2. Spaan JAE, Piek JJ, Siebes M. Coronary Circulation and Hemodynamics. Heart Physiology and Pathophysiology. Elsevier; 2001. pp. 19–44. doi:10.1016/B978-012656975-9/50004-3

3. Squires TM, Quake SR. Microfluidics: Fluid physics at the nanoliter scale.

4. Faryami A, Menkara A, Viar D, Harris CA. Testing and validation of reciprocating positive displacement pump for benchtop pulsating flow model of cerebrospinal fluid production and other physiologic systems. PLOS ONE. 2022;17: e0262372. doi:10.1371/journal.pone.0262372

5. Faryami A, Menkara A, Viar D, Harris CA. Testing and Validation of Reciprocating Positive Displacement Pump for 2 Benchtop Pulsating Flow Model of Cerebrospinal Fluid Production and Other Physiologic. 2021. doi:10.1101/2021.12.26.474197

6. Spodick DH. Normal Sinus Heart Rate: Appropriate Rate Thresholds for Sinus Tachycardia and Bradycardia. Southern Medical Journal. 1996;89: 666–667. doi:10.1097/00007611-199607000-00003

7. Li WY, Strang SE, Brown DR, Smith R, Silcox DL, Li S-G, et al. Atomoxetine changes rat’s HR response to stress from tachycardia to bradycardia via alterations in autonomic function. Autonomic Neuroscience. 2010;154: 48–53. doi:10.1016/j.autneu.2009.11.003

8. Chang C-Y, Chang R-W, Hsu S-H, Wu M-S, Cheng Y-J, Kao H-L, et al. Defects in Vascular Mechanics Due to Aging in Rats: Studies on Arterial Wave Properties from a Single Aortic Pressure Pulse. Frontiers in Physiology. 2017;8. doi:10.3389/fphys.2017.00503

9. Merrill GF, Costea DM, Sharp VA. Caffeine and Pressure Flow Autoregulation. World J Cardiovasc Dis. 2019;09: 253–266. doi:10.4236/wjcd.2019.94023

10. Riva CE, Grunwald JE, Sinclair SH, Petrig BL. Blood velocity and volumetric flow rate in human retinal vessels. Invest Ophthalmol Vis Sci. 1985;26: 1124–32.

11. Williamson TH, Harris A. Ocular blood flow measurement. British Journal of Ophthalmology. BMJ Publishing Group; 1994. pp. 939–945. doi:10.1136/bjo.78.12.939

12. Rubinstein EH, Sessler DI. Skin-surface temperature gradients correlate with fingertip blood flow in humans. Anesthesiology. 1990;73: 541–5.

13. Feke GT, Tagawa H, Deupree DM, Goger DG, Sebog J, Weirer JJ. Blood Flow In the Normal Human Retina. Investigative Ophthalmology & Visual Science. 1989.

14. Maleki N, Dai W, Alsop DC. Blood flow quantification of the human retina with MRI. NMR in Biomedicine. 2011;24: 104–111. doi:10.1002/nbm.1564

15. Kaufman S, Levasseur J. Effect of portal hypertension on splenic blood flow, intrasplenic extravasation and systemic blood pressure. 2003. doi:10.1152/ajpregu.00516.2002.-We

16. Claridge KG, James CB. Ocular pulse measurements to assess pulsatile blood flow in carotid artery disease. British Journal of Ophthalmology. 1994, 78: 321–323. doi:10.1136/bjo.78.4.321

17. Langham ME, Farrell RA, O’Brien V, Silver DM, Schilder P. Blood flow in the human eye. Acta Ophthalmologica. 2009;67: 9–13. doi:10.1111/j.1755-3768.1989.tb07080.x

18. Zarrinkoob L, Ambarki K, Wåhlin A, Birgander R, Eklund A, Malm J. Blood flow distribution in cerebral arteries. Journal of Cerebral Blood Flow and Metabolism. 2015;35: 648–654. doi:10.1038/jcbfm.2014.241

19. Rebhan J, Parker LP, Kelsey LJ, Chen FK, Doyle BJ. A computational framework to investigate retinal haemodynamics and tissue stress. Biomechanics and Modeling in Mechanobiology. 2019;18: 1745–1757. doi:10.1007/s10237-019-01172-y

20. Simon AC, Levenson J. Abnormal wall shear conditions in the brachial artery of hypertensive patients. Journal of Hypertension. 1990;8: 109–114. doi:10.1097/00004872-199002000-00003

21. Irace C, Carallo C, Crescenzo A, Motti C, de Franceschi MS, Mattioli PL, et al. NIDDM is associated with lower wall shear stress of the common carotid artery. Diabetes. 1999;48: 193–197. doi:10.2337/diabetes.48.1.193

22. Fontainhas AM, Townes-Anderson E. RhoA Inactivation Prevents Photoreceptor Axon Retraction in an In Vitro Model of Acute Retinal Detachment. Investigative Opthalmology & Visual Science. 2011;52: 579. doi:10.1167/iovs.10-5744

23. Potic J, Mbefo M, Berger A, Nicolas M, Wanner D, Kostic C, et al. An in vitro Model of Human Retinal Detachment Reveals Successive Death Pathway Activations. Frontiers in Neuroscience. 2020;14. doi:10.3389/fnins.2020.571293

24. Maloca PM, Tufail A, Hasler PW, Rothenbuehler S, Egan C, Ramos de Carvalho JE, et al. 3D printing of the choroidal vessels and tumours based on optical coherence tomography. Acta Ophthalmologica. 2019;97. doi:10.1111/aos.13637

25. Tsai C-C, Kau H-C, Kao S-C, Lin M-W, Hsu W-M, Liu J-H, et al. Pulsatile ocular blood flow in patients with Graves’ ophthalmopathy. Eye. 2005;19: 159–162. doi:10.1038/sj.eye.6701434

26. Liu X, Wang X, Zhang L, Sun L, Wang H, Zhao H, et al. A novel method for generating 3D constructs with branched vascular networks 2 using multi-materials bioprinting and direct surgical anastomosis 3 4. doi:10.1101/2021.03.21.436268

27. Baüerle FK, Karpitschka S, Alim K. Living System Adapts Harmonics of Peristaltic Wave for Cost-Efficient Optimization of Pumping Performance. Physical Review Letters. 2020;124. doi:10.1103/PhysRevLett.124.098102

28. Levesque MJ, Nerem RM, Sprague EA. Vascular endothelial cell proliferation in culture and the influence of flow.

29. Khodadadei F, Liu AP, Harris CA. A high-resolution real-time quantification of astrocyte cytokine secretion under shear stress for investigating hydrocephalus shunt failure. Communications Biology. 2021;4: 387. doi:10.1038/s42003-021-01888-7

30. Moncrief K, Kaufman S. Splenic baroreceptors control splenic afferent nerve activity. Am J Physiol Regul Integr Comp Physiol. 2006;290: 352–356. doi:10.1152/ajpregu.00489.2005.-Stenosis

31. Sherwood CP, Elkington DC, Dickinson MR, Belcher WJ, Dastoor PC, Feron K, et al. Organic Semiconductors for Optically Triggered Neural Interfacing: The Impact of Device Architecture in Determining Response Magnitude and Polarity. IEEE Journal of Selected Topics in Quantum Electronics. 2021;27: 1–12. doi:10.1109/JSTQE.2021.3051408

32. Omoti AE, Omoti CE. Ocular toxicity of systemic anticancer chemotherapy. Pharm Pract (Granada). 2006;4: 55–9.

33. Al-Tweigeri T, Nabholtz J-M, Mackey JR. Ocular toxicity and cancer chemotherapy: A review. Cancer. 1996;78: 1359–1373. doi:10.1002/(SICI)1097-0142(19961001)78:7<1359::AID-CNCR1>3.0.CO;2-G

34. Steinle JJ, Zhang Q, Thompson KE, Toutounchian J, Yates CR, Soderland C, et al. Intra-Ophthalmic Artery Chemotherapy Triggers Vascular Toxicity through Endothelial Cell Inflammation and Leukostasis. Investigative Opthalmology & Visual Science. 2012;53: 2439. doi:10.1167/iovs.12-9466

